# Psychological resilience correlates with EEG source-space brain network flexibility

**DOI:** 10.1101/437848

**Authors:** Veronique Paban, Julien Modolo, Ahmad Mheich, Mahmoud Hassan

## Abstract

**Objective:** We aimed at identifying the potential relationship between the dynamical properties of the human functional network at rest and one of the most prominent traits of personality, namely resilience.

**Approach:** To tackle this issue, we used resting-state EEG data recorded from 45 healthy subjects. Resilience was quantified using the 10-items Connor-Davidson Resilience Scale (CD-RISC). Using a sliding windows approach, brain networks in each EEG frequency band (delta, theta, alpha and beta) were constructed using the EEG source-space connectivity method. Brain networks dynamics were evaluated using the network flexibility, linked with the tendency of a given node to change its modular affiliation over time.

**Main Results:** The results revealed a negative correlation between the psychological resilience and the brain network flexibility for a limited number of brain regions within the delta, alpha and beta bands.

**Significance:** This study provides evidence that network flexibility, a metric of dynamic functional networks, is strongly correlated with psychological resilience as assessed from personality testing. Beyond this proof-of-principle that reliable EEG-based quantities representative of personality traits can be identified, this motivates further investigation regarding the full spectrum of personality aspects and their relationship with functional networks.

## Introduction

An evolving field of neuroscience aims to reveal the neural substrates of personality, referring to the relatively stable character of an individual that influences her/his long-term behavioral style (Dubois et al., 2018). A key personality character is *resilience*, defined as the ability to adapt to stress, adversity, negative events and cope actively with life challenges (Fletcher and Sarkar, 2013; Luthar, 2003; Rutter, 2006). Recently, a multi-system model of resilience has been proposed by (Liu et al., 2017), in which resilience comprises three structures: (1) the innermost layer comprises the physiological, biological and demographic profiles of an individual, (2) the intermediate layer includes internal factors such as family, friends, and personal experiences, and (3) the outermost layer corresponds to external resilience such as access to healthcare and social services. However, except some few efforts (Kong et al., 2015; Reynaud et al., 2013; Waugh and Koster, 2015), the neural subtracts of this complex personality trait remain unclear.

The last decade has witnessed an increase of studies that consider the human brain as a large-scale network. It is thus unsurprising that network neuroscience, which uses tools from graph theory to better understand neural systems (Bassett and Sporns, 2017), has become one of the most promising approaches to link behavior to brain function, including personality traits (Markett et al., 2018). In the network neuroscience model, the human brain is summarized by a set of nodes representing brain regions and a set of edges representing the connections between these brain regions. The magneto/electro-encephalography (M/EEG) source-space networks provide a unique direct and non-invasive access to those electrophysiological brain networks, at the milliseconds temporal scale (Hassan and Wendling, 2018; Mheich et al., 2018). The excellent time precision of this method allows the tracking of the brain networks dynamics at unprecedented time scale. In this paper, we aimed at test the hypothesis that metrics derived from brain networks dynamics can be correlated to resilience scores.

To test this hypothesis, we used resting-state EEG data recorded from N=45 healthy subjects. Resilience was quantified using the Connor-Davidson Resilience Scale (CD-RISC) (Campbell-Sills and Stein, 2007), where higher scores correspond to greater resilience. Brain networks in each frequency band (delta, theta, alpha and beta) were constructed using the EEG source-space connectivity method. By applying multislice modularity algorithms on the dynamic networks, the reconfiguration of EEG-source space networks was quantified using flexibility, defined as how often a given node changes its modular affiliation over time, computed at the level of each brain region. Results revealed essentially a negative correlation between psychological resilience and EEG-based brain functional network flexibility.

## Material and methods

### Participants

A total of forty-five healthy subjects were recruited (22 women). The mean age was 34.7 years old (SD=9.1 years, range=18-55). Education ranged from 10 years of schooling to the Ph.D. degree. None of the volunteers reported taking any medication or drugs nor suffering from any past or present neurological or psychiatric disease. After receiving approval from the Aix-Marseille university ethics committee according to the Declaration of Helsinki, participants filled out the CD-RISC questionnaire at home approximately 1 week before the EEG experiment. Written informed consent was obtained from all participants prior to study onset.

### Measuring Psychological Resilience

The Connor-Davidson Resilience Scale (CD-RISC) (Connor and Davidson, 2003) is a 25-item scale that measures the ability to cope with adversity. A 10 items version (CD-RISC 10) of this scale has been developed by Campbell-Sills and Stein (Campbell-Sills and Stein, 2007). A 10 items version validated for French speaking populations was used in the present study (Guihard et al., 2018; Scali et al., 2012). The 10 items are rated on a five-point Likert scale that ranges from 0 (not at all) to 4 (true nearly all of the time). Higher scores correspond to greater resilience. This scale demonstrated good internal consistency and construct validity (Campbell-Sills and Stein, 2007). In our sample, the CD-RISC 10 exhibited a reliability of α = 0.90.

### Data acquisition and preprocessing

Each EEG session consisted in a 10-min resting period with the participant’s eyes closed (Paban et al., 2018). Participants were seated in a dimly lit room, were instructed to close their eyes and then to simply relax until they were informed that they could open their eyes. Participants were instructed that the resting period would last approximately 10 min. The eyes-closed resting EEG recordings protocol was chosen to minimize movement and sensory input effects on electrical brain activity. EEG data were collected using a 64-channel Biosemi ActiveTwo system (Biosemi Instruments, Amsterdam, The Netherlands) positioned according to the standard 10-20 system montage, 1 electrocardiogram, and 2 bilateral electro-oculogram electrodes (EOG) for horizontal movements. Nasion-inion and preauricular anatomical measurements were made to locate each individual’s vertex site. Electrode impedances were kept below 20 kOhm. EEG signals are frequently contaminated by several sources of artifacts, which were addressed using the same preprocessing steps as described in several previous studies dealing with EEG resting-state data (Kabbara et al., 2018; Kabbara et al., 2017; Rizkallah et al., 2018). Briefly, bad channels (signals that are either completely flat or contaminated by movement artifacts) were first identified by visual inspection, complemented by the power spectral density. These bad channels were then recovered using an interpolation procedure implemented in Brainstorm (Tadel et al., 2011), using neighboring electrodes within a 5cm radius. Epochs with voltage fluctuations >+80 μV and <−80 μV were removed. Five artifact-free epochs of 40s lengths were selected for each participant. This epoch length was used in a previous study, and was considered as a good compromise between the needed temporal resolution and the results reproducibility (Kabbara et al., 2017). Using a sliding window approach to compute the functional connectivity, a large number of networks (depend on the analyzed frequency band) were obtained for each 40s-epoch.

### Brain networks construction

First, brain networks were reconstructed using the “EEG source-space connectivity” method (Hassan et al., 2014; Hassan and Wendling, 2018), which includes two main steps: 1) reconstruct the dynamics of the cortical sources by solving the inverse problem, and 2) measure the statistical couplings (functional connectivity) between the reconstructed time series. EEGs and MRI template (ICBM152) were co-registered through the identification of anatomical landmarks using Brainstorm (Tadel et al., 2011). A Desikan-Killiany atlas-based segmentation approach was used, consisting in 68 cortical regions (Desikan *et al.*, 2006). The OpenMEEG (Gramfort et al., 2010) software was used to compute the head model. Here, we used the weighted minimum norm estimate (wMNE) algorithm as an inverse solution. The reconstructed regional time series were filtered in different frequency bands [delta (1-4Hz), theta (4-8 Hz); alpha (8-13 Hz); beta (13-30 Hz)]. For each frequency band, functional connectivity was computed between the regional time series using the phase locking value (PLV) measure (Lachaux *et al.*, 1999). This combination wMNE/PLV was chosen according to a recent model-based comparative study of different inverse/connectivity combinations (Hassan et al., 2017) Using PLV, dynamic functional connectivity matrices were computed for each epoch using a sliding window technique. It consists in moving a time window of certain duration *δ*, and PLV is calculated within each window. As recommended in (Lachaux et al., 2000), we selected the smallest window length equal to 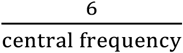 where 6 is the number of ‘cycles’ at the given frequency band. For instance, in the theta band, since the central frequency (Cf) equals to 6 Hz, *δ* equals 1s, and *δ*=279 ms (Cf=21.5 Hz) in beta band. Thus, for each epoch, 33 networks were obtained for the theta band, and 130 networks in the beta band. The same calculation was adopted for other frequency bands. Finally, we kept only the strongest 10% of connections.

### Network modularity and flexibility

The obtained dynamic matrices were divided into time-dependent modules using the multislice community detection approach described in (Mucha et al., 2010). It consists in introducing a parameter that associates nodes across time, before applying the modularity procedure and is defined as:

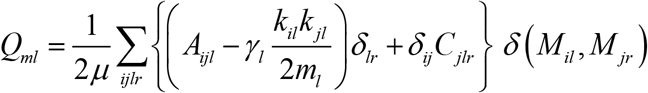

where nodes *i* and *j* are assigned to modules *M*_*il*_ and *M*_*jl*_ in window *l*, respectively. *A*_*ijl*_ represents the weight of the edge between these two nodes, and *γ*_*l*_ is the structural resolution parameter of layer *l*. *C*_*jlr*_ is the connection strength between node *j* in layer *r* and node *j* in layer *l*. In this work, the structural resolution parameter *γ* and the inter-layer coupling parameter *C* were set to 1. *k*_*il*_ is the strength of node *i* in layer *l*, the *δ*-function *δ(x, y)* is 1 if *x* = *y* and 0 otherwise,

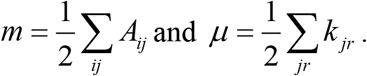

The multilayer network modularity was computed 100 times since Q may vary from run to run (degeneracy). We produced a consensus matrix whose elements indicate the ratio of each region to be in the same module with the other regions among these 100 partitions. Only elements in the consensus matrix higher than an appropriate random null model were considered. To quantify the dynamics of brain networks, we used the flexibility metric proposed in (Bassett et al., 2011). The flexibility of a brain region is defined as the number of times that a brain region changed modular assignment throughout the session, normalized by the total number of changes that were possible.

### Statistical analysis

Since data were not normally distributed, we assessed the statistical difference between the two groups using the Mann Whitney U Test, also known as Rank-Sum Wilcoxon test. To deal with the family-wise error rate, the statistical tests were corrected for multiple comparisons (N=68) using the False Discovery Rate (FDR) method.

### Software

The functional connectivity, network measures and network visualization were performed using BCT (Rubinov and Sporns, 2010), EEGNET (Hassan et al., 2015) and BrainNet viewer (Xia et al., 2013), respectively. The Network Community Toolbox (http://commdetect.weebly.com/) was used to compute the consensus matrices as well as the values provided by the flexibility metrics.

## Results

The study is summarized in Figure 1. First, EEG data were recorded and preprocessed. Second, the dynamic networks were estimated using the EEG source-space approach giving a set of brain network at the given time period, for each frequency band. Third, the flexibility of each brain region in each frequency bands were computed for each subject. Finally, the correlation between the brain regions flexibility and the resilience score (CD-RISC) was computed.

**Figure 1:**
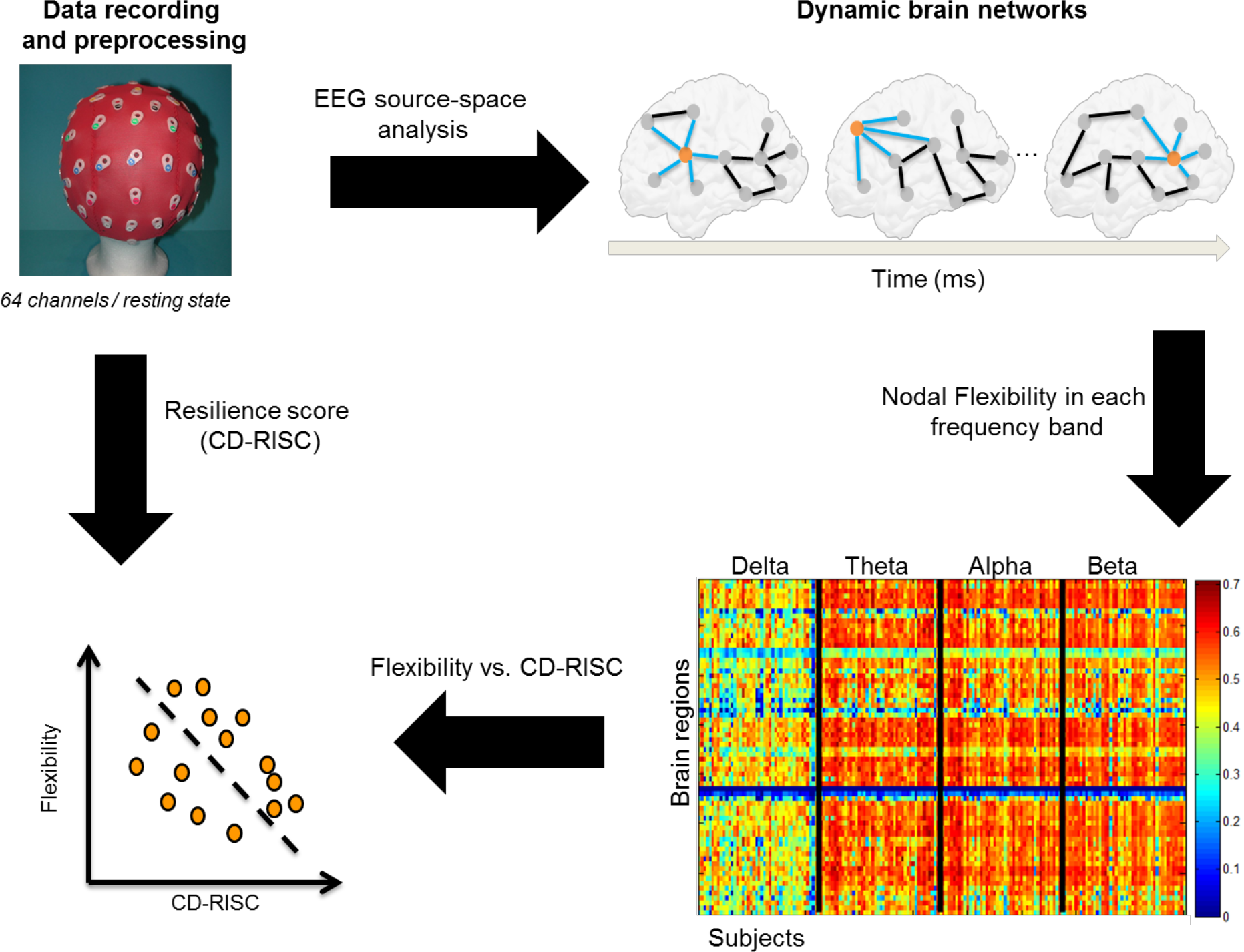
Experimental protocol and data analysis pipeline. Then, we computed the Pearson correlation between the resilience score and the global flexibility (averaged over all brain regions for each subject in each frequency band). The corresponding results are presented in Figure 2. For all frequency bands, a negative correlation was observed. This negative correlation was significant for delta (*R*=*−0.51, p*=*0.0003*), alpha (*R*=*−0.41, p*=*0.004*) and beta (*R*=*−0.43, p*=*0.002*) bands, while non-significant for the theta band (*R*=*−0.19, p*=*0.2*).

**Figure 2.**
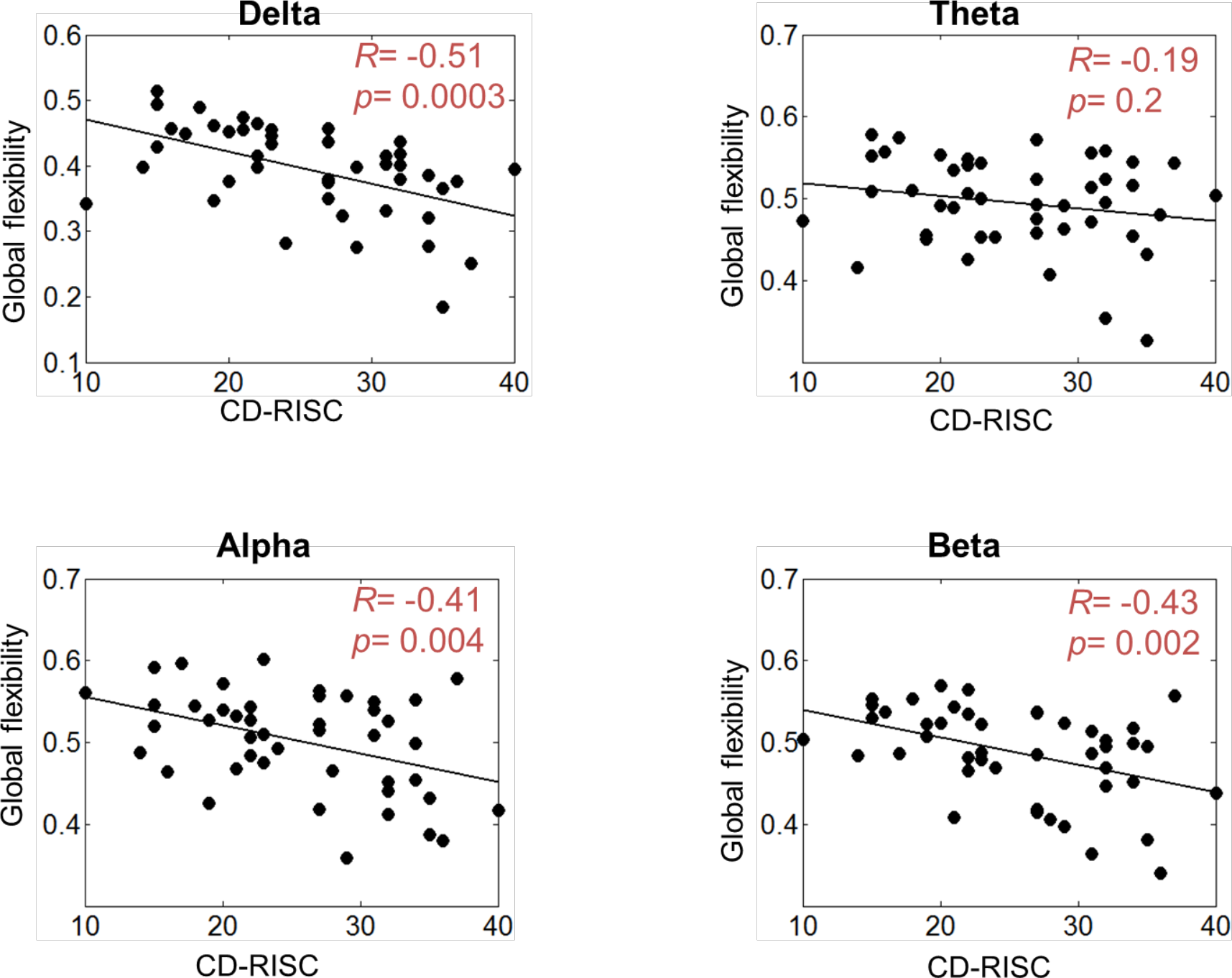
Correlation between the global flexibility (averaged over all brain regions) and the CD-RISC score. We then focused on the correlation between flexibility and the resilience score at the level of each brain region. In the delta band, brain regions that were significant correlated with the CD-SCORE are illustrated in Figure 3. The flexibility of the left cuneus (*R*=*−0.52, p*=*0.0002, FDR corrected*), the right cuneus (*R*=*−0.50, p,*=*0.0004, FDR corrected*), the left superior parietal (*R*=*−0.49, p*=*0.0005, FDR corrected*), the right superior parietal (*R*=*−0.49, p*=*0.0006, FDR corrected*), and the right entorhinal (*R*=*−0.45, p*=*0.0006, FDR corrected*), had the highest (> 90%) correlation with the resilience score.

**Figure 3.**
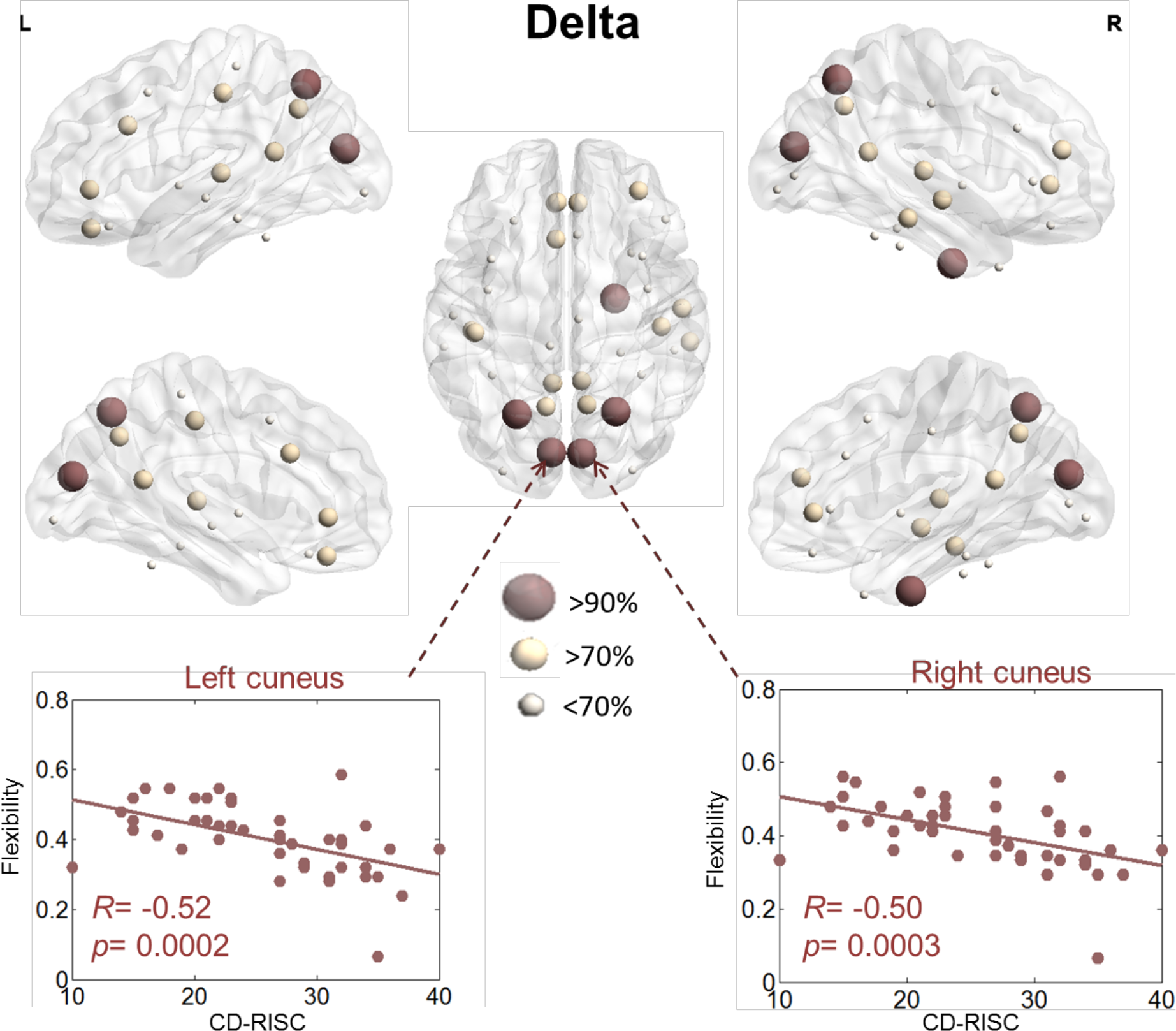
Illustration of the brain regions for which flexibility is significantly correlated with resilience within the delta band. The results for the alpha band are presented in Figure 4. The flexibility of the left caudal anterior cingulate (*R*=*−0.46, p*=*0.001, FDR corrected*), the right rostral middle frontal (*R*=*−0.46, p*=*0.001, FDR corrected*), the right pars orbitalis (*R*=*−0.45, p*=*0.001, FDR corrected*), the left inferior parietal (*R*=*-0.44, p*=*0.002, FDR corrected*), the left isthmus cingulate (*R*=*−0.43, p*=*0.003, FDR corrected*), the right isthmus cingulate (*R*=*−0.42, p*=*0.004, FDR corrected*), and the right pars opercularis (*R*=*−0.41, p*=*0.004, FDR corrected*), showed the highest correlations with the resilience score.

**Figure 4.**
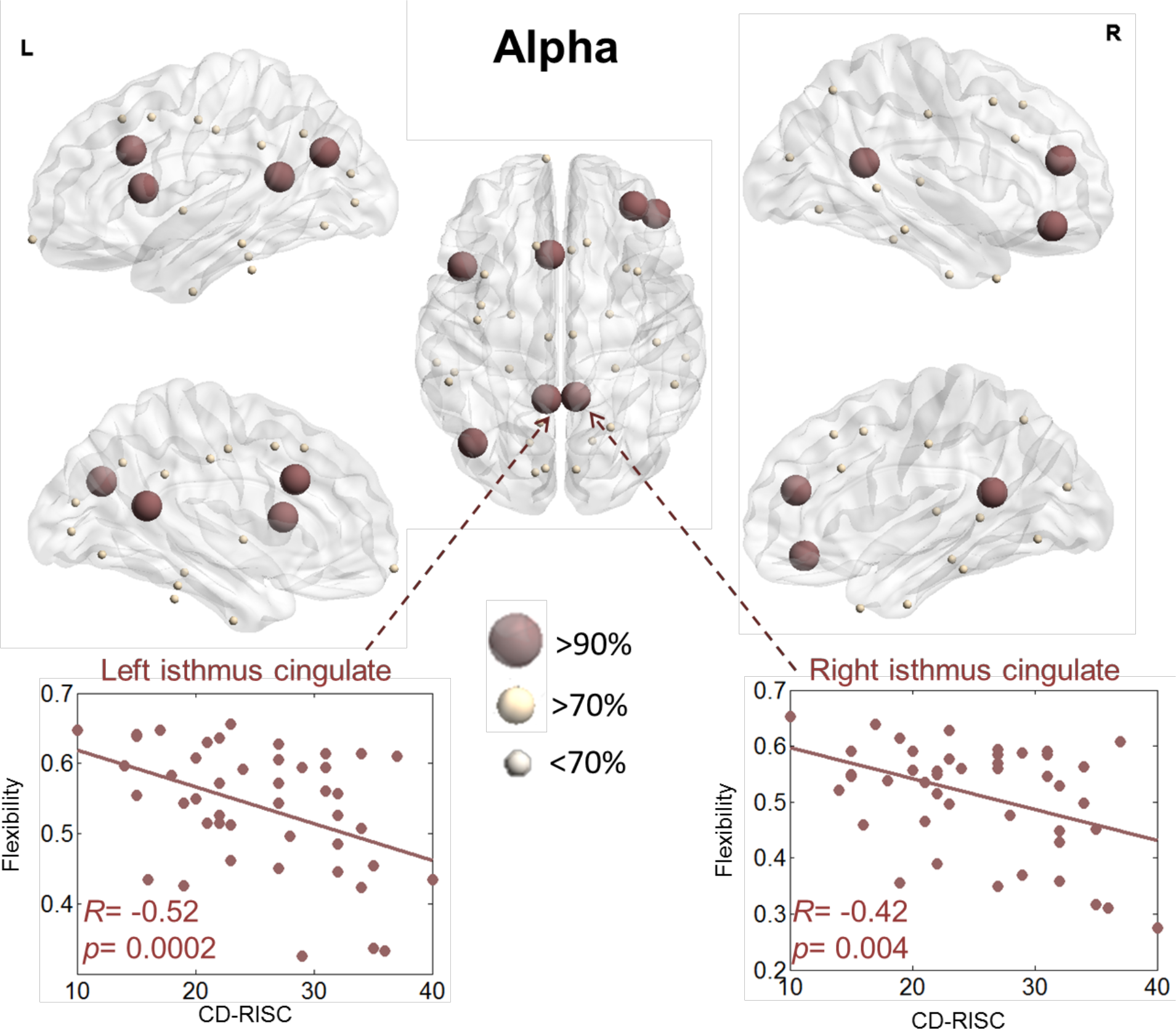
Illustration of the brain regions for which flexibility is significantly correlated with resilience within the alpha band. Results regarding the beta band revealed a more pronounced implication of visual networks. Specifically, the flexibility of the left lingual (*R*=*-0.50, p*=*0.0004, FDR corrected*), right lingual (*R*=*-0.48, p*=*0.0007, FDR corrected*), left pericalcarine (*R*=*−0.48, p*=*0.0007, FDR corrected*), right pericalcarine (*R*=*−0.47, p*=*0.0009, FDR corrected*), left cuneus (*R*=*−0.46, p*=*0.001, FDR corrected*), left isthmus cingulate (*R*=*−0.46, p*=*0.001, FDR corrected*), left medial orbitofrontal (*R*=*−0.45, p*=*0.001, FDR corrected*), and left lateral orbitofrontal (*R*=*−0.45, p*=*0.001, FDR corrected*) had the highest correlation with the resilience score.

**Figure 5.**
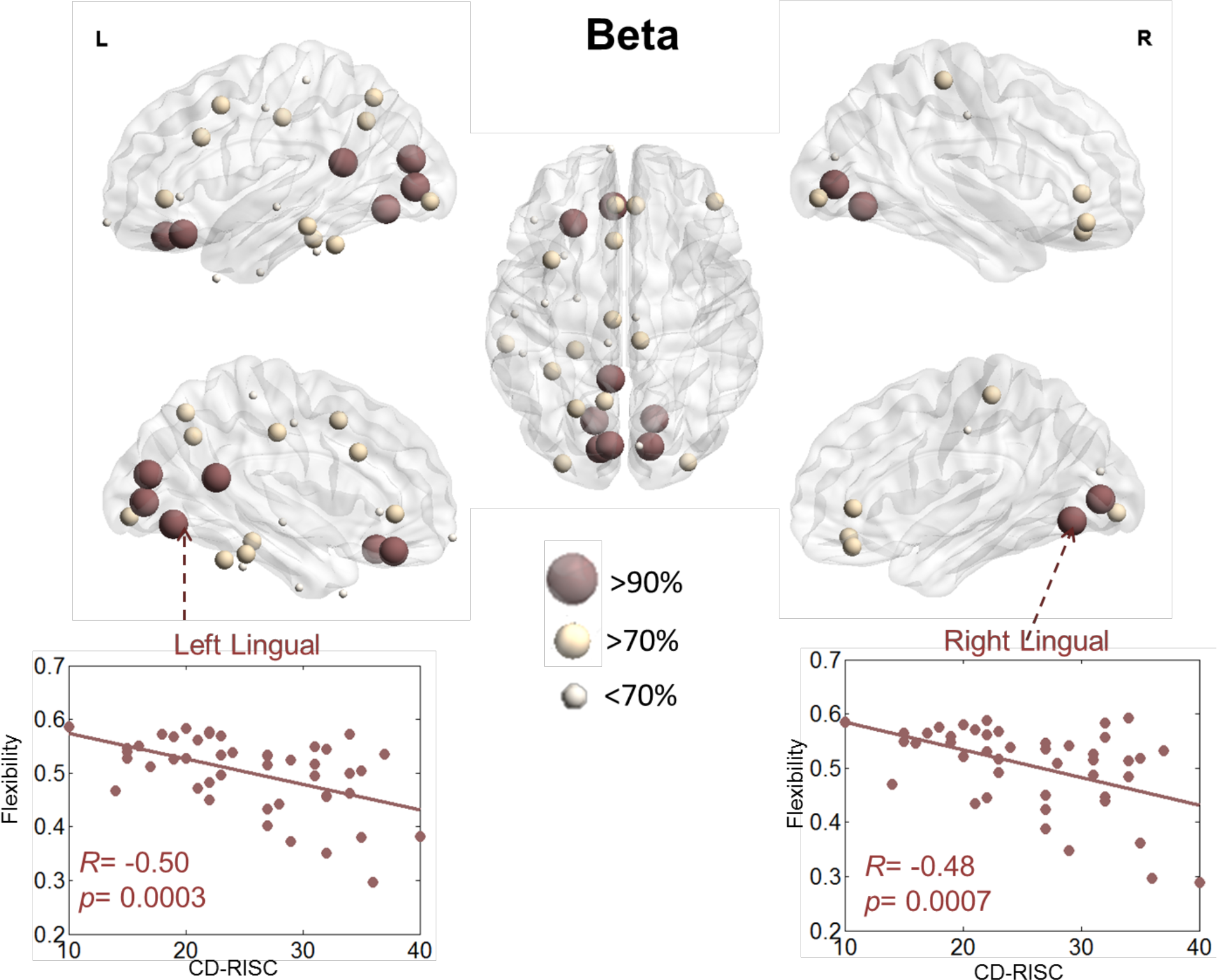
Illustration of the brain regions for which flexibility is significantly correlated with resilience within the alpha band.

## Discussion

Our results reveal a robust and direct relationship between the network flexibility (as quantified through graph theory) of human functional brain networks and resilience (as measured with neuropsychological testing). This result has been obtained using non-invasive brain recordings (EEG), in the absence of any specific task or stimulus (resting-state recordings), for a relatively large sample size (N=45). Specifically, the network flexibility of specific brain regions is shown to be significantly decreased when the resilience score is higher, which can appear non-intuitive: if resilience refers to a capacity to cope with external stress and adapt, it could imply that brain networks can reconfigure to adapt to these changes. Our results point at the opposite: most brain regions negatively correlated with the resilience score are part of the “rich club” (“core” functional network of brain regions (Van Den Heuvel and Sporns, 2011)) and illustrate that the brain core network is less flexible in resilient subjects. This suggests that a stable core network would allow preventing excessive reconfiguration of functional brain networks following external stress factors for example. From a fundamental perspective, these results also suggest that there exists a brain network of critical importance, which underlies one of the most prominent personality traits (resilience). This raises the possibility to probe further potential brain network correlates of other personality traits in the future. Furthermore, this link between core network structures and resilience is fundamental, since it can be detected non-invasively even in resting-state recordings. Another insight from our analysis is that dynamic, but not static, network analysis was able to reveal this association between resilience and a graph theory metric. Static analyses were indeed performed using clustering coefficient, participation coefficient, strength and efficiency, and did not reveal any significant correlation with the resilience score (results not shown). Therefore, this highlights the importance of the dynamic reconfiguration of brain networks in shaping behavior. Recent studies have emphasized the interest to characterize the architecture underlying these dynamics, for example by identifying the “life duration” and transition rates of these frequency-specific networks (Vidaurre et al., 2018). Overall, as emphasized in a recent perspective paper Tompson et al. (Tompson et al., 2018), there is a significant potential regarding the use of network neuroscience tools to unveil the brain networks underlying personality traits, with flexibility and integration being especially promising.

### Neural substrates of resilience

Few functional neuroimaging studies have examined brain networks associated with resilience. Most of them mainly focused on patient populations, such as depressive, traumatized and post-traumatic stress disorders, in which alterations were described in regions of the brain involved in emotion and stress regulation circuitry, see (van der Werff et al., 2013) for review. In healthy adults, to our knowledge, only one resting-state fMRI (rs-fMRI) experiment has been reported to date. Interestingly, Kong et al. reported that psychological resilience had significant negative correlations with the rs-fMRI signals, but in regions such as the bilateral insula, right dorsal and rostral anterior cingulate cortex (Kong et al., 2015). Our study did not highlight the same brain regions, since our dynamic network analysis identified regions belonging to the “core” functional network described by (Van Den Heuvel and Sporns, 2011). Those regions are supposed to play a central role in establishing and maintaining efficient global brain communication. Core regions are involved in cognitive processes related to top-down attentional control (superior parietal cortex (Sestieri et al., 2017)), decision making (subdivision of the orbitofrontal cortex; (Schuck et al., 2018)), cognitive regulation of behavior (caudal anterior cingulate; (Bush et al., 2000)), and spatial location (entorhinal cortex; (Kim and Maguire, 2018)). Our data also highlighted regions participating in the visual network (e.g. cuneus, the lingual gyrus, and the pericalcarine cortex). This network was present in the beta frequency band, which is thought to reflect different aspects of sensory information processing (Hong et al., 2008). Another region negatively correlated with the resilience score is the inferior parietal cortex, which has been shown to be involved in major cognitive functions, including visuo-spatial attention and visual memory (Egner et al., 2008).

The present study provided evidence of neuronal substrates of resilience. In healthy adults, it makes sense that the ability of individual to cope actively with life’s challenge (Fletcher and Sarkar, 2013) requires brain regions involved in high-level cognitive processes. To be efficient, these regions should have a modular organization stable over time. As suggested by (Betzel et al., 2016), such network architecture limits information exchange across modules, which allows a specialized information processing. Such modular organization seemed to be an important factor to maintain a stable equilibrium of psychological and physical functioning when facing adversity, ranging from daily problems to major life events (Bonanno, 2004). When this is not the case, i.e., when few brain regions showed high modular affiliation exchanges, one may assume that they multiply their contributions and so participate in a multitude of processes. In this context, subjects had low-resilient scores, meaning that they exhibit passive coping strategies, high emotional load, ruminative and depressogenic thinking, and low life satisfaction (Davydov et al., 2010).

### Methodological considerations

First, in the present study, we used a template source space, instead of a subject-specific one, i.e. the same structural MRIs of healthy subjects was used in our EEG functional connectivity analysis. In the case of healthy subjects, it was reported that co-registration with the template yielded largely consistent connectivity and network estimates as compared to native MRI (Douw et al., 2018). However, in the case of patient-related studies, the method of choice is to use patient-specific MRIs.

Second, it is important to keep in mind that computing functional connectivity at the scalp level is severely corrupted by the volume conduction problem (Brookes et al., 2014; Lai et al., 2018; Schoffelen and Gross, 2009). M/EEG connectivity analysis at the source-space was shown to reduce this effect as connectivity is applied to “local” time-series generated by cortical neuronal assemblies modelled-for instance-as current dipole sources. Yet, the “mixing effects” can also occur in the source-space. To address this issue, a number of methods were developed based mostly on the rejection of zero-lag correlation. In particular, “unmixing” methods, called “leakage correction” (such as the orthogonalization approach (Colclough et al., 2015)), have been reported, which force the reconstructed signals to have zero cross-correlation at lag zero. Here we prefered to use the PLV metric and keep these zero-lag correlations, since several experimental studies reported the importance (in many conditions) of zero-lag synchronisations in the human brain. Nevertheless, we believe that there is no ideal solution yet for this methdological issue and that further efforts are needed to completely solve the spatial leakage problem.

## References

Bassett, D.S., and Sporns, O. (2017). Network neuroscience. Nature neuroscience 20, 353.

Bassett, D.S., Wymbs, N.F., Porter, M.A., Mucha, P.J., Carlson, J.M., and Grafton, S.T. (2011). Dynamic reconfiguration of human brain networks during learning. Proceedings of the National Academy of Sciences.

Betzel, R.F., Fukushima, M., He, Y., Zuo, X.-N., and Sporns, O. (2016). Dynamic fluctuations coincide with periods of high and low modularity in resting-state functional brain networks. NeuroImage 127, 287–297.

Bonanno, G.A. (2004). Loss, trauma, and human resilience: Have we underestimated the human capacity to thrive after extremely aversive events? American psychologist 59, 20.

Brookes, M.J., Woolrich, M.W., and Price, D. (2014). An Introduction to MEG connectivity measurements. In Magnetoencephalography (Springer), pp. 321–358.

Bush, G., Luu, P., and Posner, M.I. (2000). Cognitive and emotional influences in anterior cingulate cortex. Trends in cognitive sciences 4, 215–222.

Campbell-Sills, L., and Stein, M.B. (2007). Psychometric analysis and refinement of the Connor-Davidson resilience scale (CD-RISC): validation of a 10-item measure of resilience. Journal of Traumatic Stress: Official Publication of The International Society for Traumatic Stress Studies 20, 1019–1028.

Colclough, G., Brookes, M., Smith, S., and Woolrich, M. (2015). A symmetric multivariate leakage correction for MEG connectomes. NeuroImage 117, 439–448.

Connor, K.M., and Davidson, J.R. (2003). Development of a new resilience scale: The Connor-Davidson resilience scale (CD-RISC). Depression and anxiety 18, 76–82.

Davydov, D.M., Stewart, R., Ritchie, K., and Chaudieu, I. (2010). Resilience and mental health. Clinical psychology review 30, 479–495.

Douw, L., Nieboer, D., Stam, C.J., Tewarie, P., and Hillebrand, A. (2018). Consistency of magnetoencephalographic functional connectivity and network reconstruction using a template versus native MRI for co-registration. Hum Brain Mapp 39, 104–119.

Dubois, J., Galdi, P., Han, Y., Paul, L.K., and Adolphs, R. (2018). Resting-state functional brain connectivity best predicts the personality dimension of openness to experience.

Egner, T., Monti, J.M., Trittschuh, E.H., Wieneke, C.A., Hirsch, J., and Mesulam, M.-M. (2008). Neural integration of top-down spatial and feature-based information in visual search. Journal of Neuroscience 28, 6141–6151.

Fletcher, D., and Sarkar, M. (2013). Psychological resilience: A review and critique of definitions, concepts, and theory. European Psychologist 18, 12.

Gramfort, A., Papadopoulo, T., Olivi, E., and Clerc, M. (2010). OpenMEEG: opensource software for quasistatic bioelectromagnetics. Biomedical engineering online 9, 45.

Guihard, G., Deumier, L., Alliot-Licht, B., Bouton-Kelly, L., Michaut, C., and Quilliot, F. (2018). Psychometric validation of the French version of the Connor-Davidson Resilience Scale. L’Encéphale 44, 40–45.

Hassan, M., Dufor, O., Merlet, I., Berrou, C., and Wendling, F. (2014). EEG source connectivity analysis: from dense array recordings to brain networks. PloS one 9, e105041.

Hassan, M., Merlet, I., Mheich, A., Kabbara, A., Biraben, A., Nica, A., and Wendling, F. (2017). Identification of Interictal Epileptic Networks from Dense-EEG. Brain topography 30, 60–76.

Hassan, M., Shamas, M., Khalil, M., El Falou, W., and Wendling, F. (2015). EEGNET: An Open Source Tool for Analyzing and Visualizing M/EEG Connectome. PloS one 10, e0138297.

Hassan, M., and Wendling, F. (2018). Electroencephalography Source Connectivity: Aiming for High Resolution of Brain Networks in Time and Space. IEEE Signal Processing Magazine 35, 81–96.

Hong, L.E., Summerfelt, A., Mitchell, B.D., McMahon, R.P., Wonodi, I., Buchanan, R.W., and Thaker, G.K. (2008). Sensory gating endophenotype based on its neural oscillatory pattern and heritability estimate. Archives of general psychiatry 65, 1008–1016.

Kabbara, A., Eid, H., El Falou, W., Khalil, M., Wendling, F., and Hassan, M. (2018). Reduced integration and improved segregation of functional brain networks in Alzheimer’s disease. Journal of neural engineering 15, 026023.

Kabbara, A., Falou, W.E., Khalil, M., Wendling, F., and Hassan, M. (2017). The dynamic functional core network of the human brain at rest. Scientific reports 7, 2936.

Kim, M., and Maguire, E.A. (2018). Thalamus, subiculum and retrosplenial cortex encode 3D head direction information in volumetric space. bioRxiv, 335976.

Kong, F., Wang, X., Hu, S., and Liu, J. (2015). Neural correlates of psychological resilience and their relation to life satisfaction in a sample of healthy young adults. Neuroimage 123, 165–172.

Lachaux, J.-P., Rodriguez, E., Le Van Quyen, M., Lutz, A., Martinerie, J., and Varela, F.J. (2000). Studying single-trials of phase synchronous activity in the brain. International Journal of Bifurcation and Chaos 10, 2429–2439.

Lai, M., Demuru, M., Hillebrand, A., and Fraschini, M. (2018). A comparison between scalp-and source-reconstructed EEG networks. Scientific reports 8, 12269.

Liu, J.J., Reed, M., and Girard, T.A. (2017). Advancing resilience: An integrative, multi-system model of resilience. Personality and Individual Differences 111, 111–118.

Luthar, S.S. (2003). Resilience and vulnerability: Adaptation in the context of childhood adversities (Cambridge University Press).

Markett, S., Montag, C., and Reuter, M. (2018). Network neuroscience and personality. Personality Neuroscience 1.

Mheich, A., Hassan, M., Khalil, M., Gripon, V., Dufor, O., and Wendling, F. (2018). SimiNet: a novel method for quantifying brain network similarity. IEEE transactions on pattern analysis and machine intelligence 40, 2238–2249.

Mucha, P.J., Richardson, T., Macon, K., Porter, M.A., and Onnela, J.-P. (2010). Community structure in time-dependent, multiscale, and multiplex networks. science 328, 876–878.

Paban, v., Deshayes, c., Ferrer, M.-H., Weill, a., and Alescio-Lautier, B. (2018). Resting brain functional networks and trait coping. Brain connectivity.

Reynaud, E., Guedj, E., Souville, M., Trousselard, M., Zendjidjian, X., El Khoury-Malhame, M., Fakra, E., Nazarian, B., Blin, O., and Canini, F. (2013). Relationship between emotional experience and resilience: An fMRI study in fire-fighters. Neuropsychologia 51, 845–849.

Rizkallah, J., Benquet, P., Kabbara, A., Dufor, O., Wendling, F., and Hassan, M. (2018). Dynamic reshaping of functional brain networks during visual object recognition. Journal of neural engineering.

Rubinov, M., and Sporns, O. (2010). Complex network measures of brain connectivity: uses and interpretations. Neuroimage 52, 1059–1069.

Rutter, M. (2006). Implications of resilience concepts for scientific understanding. Annals of the New York Academy of Sciences 1094, 1–12.

Scali, J., Gandubert, C., Ritchie, K., Soulier, M., Ancelin, M.-L., and Chaudieu, I. (2012). Measuring resilience in adult women using the 10-items Connor-Davidson Resilience Scale (CD-RISC). Role of trauma exposure and anxiety disorders. PloS one 7, e39879.

Schoffelen, J.M., and Gross, J. (2009). Source connectivity analysis with MEG and EEG. Human brain mapping 30, 1857–1865.

Schuck, N.W., Wilson, R., and Niv, Y. (2018). A state representation for reinforcement learning and decision-making in the orbitofrontal cortex. In Goal-Directed Decision Making (Elsevier), pp. 259–278.

Sestieri, C., Shulman, G.L., and Corbetta, M. (2017). The contribution of the human posterior parietal cortex to episodic memory. Nature Reviews Neuroscience 18, 183.

Tadel, F., Baillet, S., Mosher, J.C., Pantazis, D., and Leahy, R.M. (2011). Brainstorm: a user-friendly application for MEG/EEG analysis. Computational intelligence and neuroscience 2011, 8.

Tompson, S., Falk, E.B., Vettel, J.M., and Bassett, D.S. (2018). Network Approaches to Understand Individual Differences in Brain Connectivity: Opportunities for Personality Neuroscience. Personality Neuroscience.

Van Den Heuvel, M.P., and Sporns, O. (2011). Rich-club organization of the human connectome. Journal of Neuroscience 31, 15775–15786.

van der Werff, S.J., van den Berg, S.M., Pannekoek, J.N., Elzinga, B.M., and Van Der Wee, N.J. (2013). Neuroimaging resilience to stress: a review. Frontiers in behavioral neuroscience 7, 39.

Vidaurre, D., Hunt, L.T., Quinn, A.J., Hunt, B.A., Brookes, M.J., Nobre, A.C., and Woolrich, M.W. (2018). Spontaneous cortical activity transiently organises into frequency specific phase-coupling networks. Nature communications 9, 2987.

Waugh, C.E., and Koster, E.H. (2015). A resilience framework for promoting stable remission from depression. Clinical Psychology Review 41, 49–60.

Xia, M., Wang, J., and He, Y. (2013). BrainNet Viewer: a network visualization tool for human brain connectomics. PloS one 8, e68910.

